# Task-Dependent Motor Unit Recruitment and Rate Coding Reveal Redistribution of Neural Drive in the Human Hand

**DOI:** 10.64898/2025.12.16.694568

**Authors:** Marius Osswald, Jaime Ibáñez, Martin Regensburger, Nick Unbehaun, Alessandro Del Vecchio

## Abstract

Although Henneman’s size principle dictates an orderly small-to-large activation for a given common input, evidence suggests a certain level of flexibility in the recruitment of spinal motoneurons depending on task demands. Here, we investigate motor unit (MU) recruitment flexibility in the human first dorsal interosseous (FDI) muscle while controlling for overall muscle activation across two functionally distinct tasks using high-density intramuscular EMG (HD-iEMG) electrode arrays. Six participants performed isometric index finger abduction where the FDI serves as the prime mover, and flexion with the FDI functioning as a synergist. Recruitment thresholds (RTs) and recruitment orders (ROs) were highly consistent within the same task, but differed significantly between abduction and flexion. Across participants, 45.3% of MUs showed changes in recruitment above the coefficient of repeatability across tasks compared to only 5.0% within tasks. The spatial distribution of muscle activation did not shift between tasks, confirming that the same muscle region was engaged in both. Changes in RT were accompanied by corresponding adaptations in discharge rate (DR), preserving the inverse RT-DR relationship. Intramuscular coherence analysis revealed no differences in the delta (1-5 Hz) or alpha (5-13 Hz) band, but beta band (13-30 Hz) coherence was significantly lower during flexion than abduction, indicating reduced common input synchronization in this frequency range when FDI serves as a synergist muscle. Together, these results indicate different distributions of net excitatory input to FDI MUs across different functional tasks, potentially involving a stronger contribution of spinal circuits during flexion, as suggested by the reduced intramuscular beta coherence. Moreover, the present findings also demonstrate that common synaptic input and not intrinsic motoneuron properties determines the inverse relation between MU RT and DR.

## Introduction

Muscle force generation and movement originate from commands issued by the central nervous system. These commands are transmitted through descending cortical, subcortical, and spinal pathways that converge on *α*-motoneurons in the anterior horn of the spinal cord, which transmit the neural signals to their respective innervated muscle fibers, forming individual motor units (MUs). The recruitment and derecruitment of MUs are key neural mechanisms of force modulation and are traditionally described by Henneman’s size principle, which states that MUs are recruited in an orderly fashion from the smallest to the largest (Henneman (1957); Henneman et al. (1965, 1974)), primarily determined by the intrinsic input resistance to the motoneurons. The principle constrains recruitment relative to the input a set of motoneurons shares, and does not by itself require a single fixed ordering of MUs within a whole muscle across functionally distinct tasks (Bawa et al. (2014); Hug et al. (2023)). The firing activity of recruited MUs is further described as an “onion-skin” scheme, where earlier-recruited MUs maintain systematically higher firing rates than later-recruited MUs (De Luca et al. (1982); De Luca and Hostage (2010); Masakado et al. (1995); Hu et al. (2014)). These principles provide an efficient mechanism for graded force production and precise control particularly for small force levels and are thought to minimize fatigue (Bawa et al. (2014); Conwit et al. (1999); Stein et al. (2005)). They have been supported in a variety of muscles and tasks, including slow isometric contractions (Milner-Brown et al. (1973); Thomas et al. (1986, 1987); Riek and Bawa (1992); Feiereisen et al. (1997); Scutter and Türker (1998); Conwit et al. (1999)), different contraction speeds (Desmedt and Godaux (1977); Calancie and Bawa (1985)), and changes in muscle length (Garland et al. (1996); Stotz and Bawa (2001)). However, studies have reported some variability in recruitment when considering different functional tasks in the form of task-related selective recruitment (Butler et al. (2005); Saboisky et al. (2006); Oßwald et al. (2025)) and changes in RO within groups of MUs in the non-human (Marshall et al. (2022)) and human biceps brachii (Ter Haar Romeny et al. (1982); Tax et al. (1989)) and FDI muscle (Desmedt and Godaux (1981); Thomas et al. (1986)). These studies suggest that the central nervous system can optimize MU recruitment thresholds depending on task demands. Limited by the concurrent sampling of only small numbers of MUs with single-channel intramuscular EMG electrodes as used in these studies, the neural determinants of such RO flexibility currently remain unclear. Desmedt and Godaux (1981) speculated that the altered RO they observed in FDI during index finger abduction versus flexion may arise from differences in the distribution of descending cortical drive. Notably, the FDI serves distinct mechanical and functional roles in these two movements. During abduction, it is the sole muscle with a moment arm in that direction and therefore acts as the prime mover. However, the FDI also has a flexion moment arm and functions as a synergist to the long extrinsic flexors during index finger flexion, contributing to metacarpophalangeal joint stabilization together with the first lumbrical muscle (Masquelet et al. (1986); Liss (2012); Ranney and Wells (1988)). In addition to further insight into the mechanisms of MU control, understanding task-related changes in recruitment and rate coding of MUs, particularly for hand and digit actions, has practical implications. Real-time MU identification has gained traction in myocontrol and human-machine interfaces, where reliability and flexibility of recruitment patterns are critical (Campbell et al. (2025); Farina et al. (2017); Chen et al. (2021); Capsi-Morales et al. (2025); Grison et al. (2024, 2025)).

In this study, we investigate the variability of MU recruitment, discharge rate and common input in the FDI muscle by comparing within and across isometric contractions of the index finger in abduction and flexion direction, which corresponds to two different functional roles of the FDI muscle as described above. Using state-of-the-art HD-iEMG electrode arrays, we are able to identify large populations of concurrently active MUs and track their activity across contraction directions. By matching the global level of FDI muscle activity across contractions, we can observe the recruitment activity of MUs on a larger scale and in more controlled conditions than previously possible. The increase in bandwidth of MU activity correlation measures in larger populations of identified MUs (Negro and Farina (2012)) allows us to apply intramuscular coherence analysis to characterize the frequency-specific synchronization of common input to FDI MUs across the two tasks that might underlie the observed recruitment flexibility.

## Materials and Methods

### Participants

Six healthy volunteers (mean age 28 *±* 3.8 years; 2 male, 4 female), all right-handed, participated in the study. None had a history of neurological or neuromuscular disorders and fulfilled standard inclusion and exclusion criteria in neuromuscular studies in healthy participants. The study protocol was approved by the ethics committee of the Friedrich-Alexander University Erlangen-Nürnberg (approval number 23-271-B) and was conducted in accordance with the Declaration of Helsinki. All participants provided their informed written consent prior to participation.

### Experimental Setup

#### 2D Index Finger Dynamometer

A custom-built dynamometer was developed to measure isometric forces of the index finger in all possible contraction directions. Detailed images of the dynamometer and the 2D force feedback are shown in supplementary Figure S1A. The setup is adjustable to different arm lengths, hand sizes, and handedness of participants. During the experimental protocol, the arm is positioned in the dynamometer with an elbow flexion angle of approximately 100*^◦^* and the arm pointing forward. The hand and digits are arranged in a “pointing” posture, with the palm facing medially and the index finger pointing forward, parallel to the ground. The thumb position is fixed with velcro straps to prevent effects of thumb posture on the flexion moment arm of the first dorsal interosseous (FDI) muscle during contractions (Hudson et al. (2009)).

The index finger is attached to a three-axis load cell (ME Messsysteme K3D40, nominal force 50 N) via a 3D-printed ring-shaped attachment positioned just proximal to the proximal interphalangeal (PIP) joint (Fig. 1A). To immobilize the wrist and eliminate force generation via wrist flexion or extension, the hand is secured on both the palmar and dorsal sides using two adjustable restraints. These restraints can be extended or retracted for fixation, release, and adaptation to different hand sizes.

**Figure 1:**
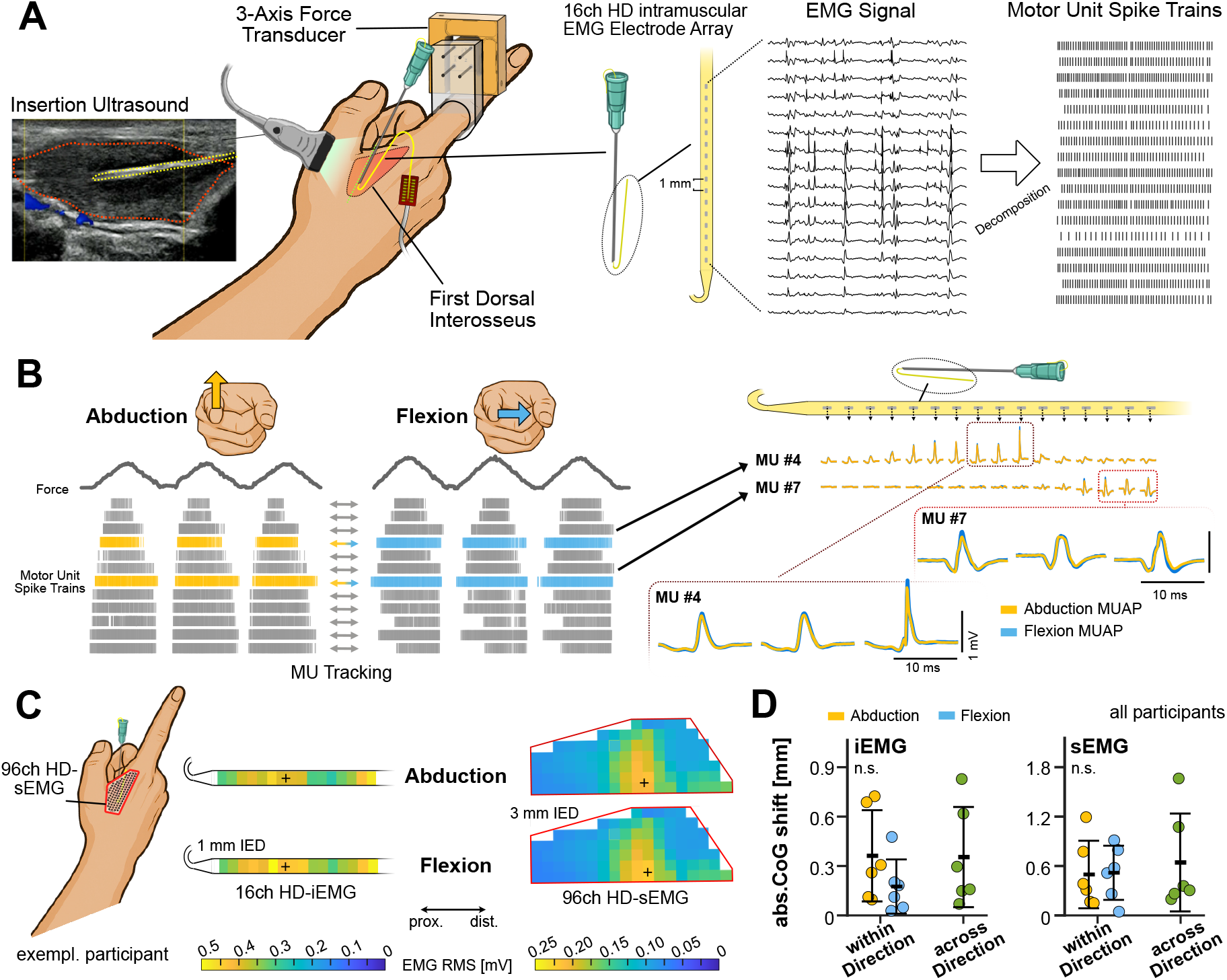
Experimental Setup and Motor Unit Tracking. **A:** Experimental setup. 16-channel high-density intramuscular electrode arrays implanted in the FDI under ultrasound guidance (red: FDI muscle volume, yellow: iEMG insertion cannula). The index finger was attached to a 3-axis load cell proximal to the PIP joint (more details see supplementary figure S1). Intramuscular EMG signals were decomposed into single motor unit activity. **B:** Motor unit tracking. Left: MUs were decomposed separately in abduction and flexion and then tracked by applying the MU filters across contraction directions. Right: Two exemplary MUs #4 and #7 are shown. MUAPs obtained from spike-triggered-averaging of each channels EMG signals shows identical MUAP shapes in both contraction directions, as well as distinct centers of activation along the electrode array. **C:** Spatial FDI activation. Both the 16ch iEMG array and a 96ch sEMG grid show highly similar spatial activity patterns for abduction and flexion. Crosses indicate the center of gravity (CoG) **D:** CoG shifts across recordings. Plots show the spatial shift in the CoGs of EMG activity amplitude for the iEMG arrays and sEMG grids in all participants. Data demonstrates the shift across contraction directions is of comparable magnitude than between trials of the same contraction direction.

The load celĺs x and y outputs are connected to an EMG amplifier system with BNC connectors. Real-time visual feedback of both force amplitude and contraction direction is provided via a custom recording interface, displaying the force vector as a dot within a circular target similar to a clock face. The distance from the center indicates force magnitude either as absolute force in N or as a percentage of maximum force. The angular position represents the contraction direction, with 12 o’clock corresponding to abduction, 9 o’clock to flexion, 6 o’clock to adduction, and 3 o’clock to extension. This mapping matched the actual orientation of the participant’s hand in the setup and can be mirrored for measurements of the left hand.

#### EMG Setup

We recorded High-Density EMG signals from the FDI muscle with multi-channel intramuscular electrode arrays designed for acute recordings (Fig. 1A) (Muceli et al. (2015); Negro et al. (2016)). The arrays contain 16 platinum-coated electrodes on a polyimide thin-film structure linearly distributed along a distance of 15 mm with an interelectrode distance of 1 mm. The electrode arrays are implanted using a 25G cannula that is removed after the insertion, with only the thin film containing the electrodes remaining in the muscle for the duration of the experimental session. In each participant, one electrode array was implanted in the FDI muscle of the right hand by a medical professional. Prior to insertion, the skin was cleaned, disinfected and the insertion point marked with a skin marker. The insertion was performed in an oblique direction from the distal third of the muscle towards its proximal third. To guarantee safe implantation, avoiding blood vessels and nerves and to ensure the placement of all 16 electrodes within muscle tissue, a desktop ultrasound device (Telemed ArtUs EXT-2H with L15-7H40-A5 probe) was used to guide the implantation in real-time (see Fig. 1A). In addition to the intramuscular electrode array, one 96-channel High-Density surface-EMG (HD-sEMG) electrode grid (GR03MM1807, OT Bioelettronica, Turin, Italy), specifically shaped to cover the full FDI muscle, was placed on the skin above the FDI. Intramuscular and surface EMG signals were recorded using a multi-channel amplifier (Quattrocento, OT Bioelettronica) at a sampling rate of 10,240 Hz with 16-bit resolution in a monopolar configuration, with the reference electrode placed on the wrist.

### Experimental Protocol

Participants were in a seated position with their arm and hand placed in the dynamometer setup as described above. As hand dominance is linked to differences in the common synaptic input to upper-limb MUs (Maillet et al. (2022)), all participants used their dominant hand. The participants performed a series of isometric contractions at constant direction of force application in index finger abduction and flexion. As the DR to MUs can have a strong influence on the estimation of coherence when comparing between conditions (Negro and Farina (2012); Negro et al. (2009); Farina et al. (2014); De la Rocha et al. (2007)), we aimed to control for comparable net neural drive to the whole FDI MU pool in both abduction and flexion by matching the global EMG amplitude of the muscle. This was achieved through a dedicated calibration protocol prior to the study protocols. The calibration protocol consisted of isometric ramp contractions at force levels of 2, 3, 4, 5, 6, 7, and 8 N in each contraction direction. For each participant, linear regression was used to determine the force levels in index finger flexion that produced the same global FDI EMG amplitude as in abduction for contractions corresponding to 10% and 20% of the participant’s maximum force in abduction. EMG amplitude was calculated as the mean root mean square (RMS) value during the plateau phase of the contractions averaged across the 16 iEMG channels. This approach of participant-specific target force selection was chosen because there is no evidence that the FDI receives similar neural drive during abduction and flexion at equivalent percentages of their respective maximum forces across individuals. In pilot data collected with the same dynamometer setup, we observed a wide spread of FDI EMG activity levels during flexion relative to abduction in contractions at matched relative force levels, as well as at matched absolute force levels. More details on the corresponding methods and results can be found in supplementary figure S2. Across participants, the calibration yielded abduction forces between 2.0 and 3.0 N and corresponding flexion forces between 1.8 and 6.2 N for the 10% maximum abduction force level to be used in the following study protocol.

The study protocol consisted of two separate parts to allow for optimized protocols for each part of the data analysis. Recruitment protocol: Participants performed a series of six isometric ramp contractions for each force level in abduction and flexion with plateau phases of 3s per contraction. Each combination of force level and contraction direction was separately recorded, from here on referred to as “trial”. In both contraction directions, concentric and eccentric phases of the ramps had durations of 5 and 10 seconds for the two force levels respectively, to ensure for equal FDI EMG RMS/s increase and decrease across force levels and direction. Neural input protocol: Participants performed two isometric ramp contractions for each force level in abduction and flexion with plateau durations of 20s per contraction and concentric and eccentric phases of 2.5s and 5s for low and high force levels. In both protocols, each trial was repeated once, resulting in two recordings with identical parameters. During data analysis only the repetition with the better initial decomposition result and sufficient force tracking accuracy was used for each trial.

The 2D real-time force feedback visualizing the force amplitude and exact contraction direction proved to be challenging for first-time users. Consequently, each participant underwent a familiarization session in the days prior to their experimental session, where they practiced isometric ramp contractions at different force amplitudes and contraction directions until sufficient precision in amplitude and direction was achieved. An initial analysis of the study data confirmed high directional accuracy of the applied force across participants during both protocols (Supplementary figure S1B).

#### Signal Processing

##### Spatial distribution of FDI muscle activation across contraction direction

To test whether abduction and flexion engaged spatially distinct regions of the FDI, we compared the spatial distribution of EMG amplitude between the two contraction directions for both the high-density intramuscular array and a high-density surface EMG grid. The data from the neural input protocol was used for this analysis.

For each contraction, the amplitude of each channel was quantified as the root-mean-square (RMS) computed over a 0.6 s maximally-overlapping window and averaged across the plateau. sEMG Channels identified as outliers were excluded on a per-participant basis. The resulting per-channel RMS values were arranged into the physical electrode layout to obtain a two-dimensional activation map.

The center of gravity (CoG) was defined as the RMS-amplitude-weighted spatial centroid of the activation map in mm. For each participant and modality, we computed the CoG separately for abduction and flexion, and quantified the CoG shift as the Euclidean distance between the two centroids. To provide a reference for the expected measurement variability, we additionally computed a within-direction shift as the CoG distance between the two repetitions of the same contraction direction, and a between-direction shift as the CoG distance between the abduction and flexion contractions. If abduction and flexion recruited the same muscle territory, the between-direction shift should not exceed the within-direction shift. For each participant the within-direction shift (averaged over abduction and flexion) and the between-direction shift were compared with a two-sided Wilcoxon signed-rank test, performed separately for the intramuscular and surface arrays in all six participants.

##### Motor Unit Identification & Tracking

The monopolar EMG signals were bandpass filtered between 100 and 4400 Hz and decomposed into single MU activity through convolutive blind source separation (Holobar and Zazula (2007); Negro et al. (2016)) using the MUedit tool (Avrillon et al. (2024a)). Relevant decomposition parameters were: decomposition iterations = 100; Extension factor = 40; Silhouette threshold = 0.9; Peel-off = On; CoV of DR threshold = 0.6; contrast function = logcosh. Subsequent visual inspection and manual editing was performed to identify and correct missed or falsely identified discharges (Del Vecchio et al. (2020)). For the data obtained from the recruitment protocol, particular attention was given to accurate identification of the first firing in each contraction, as this is critical for determining the corresponding recruitment threshold. If the first firing of a MU for a particular contraction could not be identified reliably, all identified firings in that contraction were consequently discarded.

As all contraction levels and directions were decomposed separately, MUs were tracked across force levels within the same contraction direction by concatenating the decomposed signals and application of the MU filters obtained from the decomposition from the lower onto the higher contraction level signals, and vice versa. This approach was described in previous studies with high-density surface EMG data (Frančič and Holobar (2021); Avrillon et al. (2024b)) and proved highly effective in our data (Fig. 1B). The likely reason is that intramuscular EMG provides higher temporal and spatial resolution with less low-pass filtering of the signals by volume conduction through muscle and other tissue. This increases the uniqueness and therefore separability of the action potential shapes compared to MUs data obtained from surface EMG signals. To further validate this approach and quantify the similarity of tracked MÚs MUAPs, we computed the *R*^2^ values of motor unit action potential (MUAP) shapes between abduction and flexion of tracked MUs against a null-distribution of not-tracked MUs. After removal of resulting duplicate MUs in cases where the same MU was originally identified in both contraction levels, this usually resulted in the identification of additional MUs in the higher contraction levels. This reflects the fact that the identification of low-threshold MUs is facilitated at lower contraction levels where the reduced overall neural drive results in fewer simultaneously active MUs and, consequently, less signal superposition compared to higher contraction levels. Following MU tracking across force levels, the same procedure was applied between contraction directions by applying the MU filters from the abduction to the flexion data and vice versa. The entire process was performed separately for the recruitment protocol and the coherence protocol data.

##### Motor Unit Recruitment Analysis

For the analysis of RTs and the consistency of RO between trials and contraction directions, data from the recruitment sub-protocol were used. Prior to analysis, intramuscular EMG data was band-pass filtered between 100 and 4400 Hz. Notably, removal of outlier channels was not necessary, as we did not observe any faulty or noisy signals in the data recorded in this study. EMG amplitude was quantified as the root-mean-square (RMS) calculated in windows of 400 ms with 300 ms overlap, averaged across all 16 channels of the thin-film electrode arrays. Recruitment thresholds were quantified in global EMG amplitude in mV, as this provides a more direct measure of net neural drive than absolute force (N) or relative force (% of maximum force) when considering different contraction directions. The rationale for this choice was twofold: first, motor unit recruitment is driven by increases in neural drive, with force changes representing a downstream consequence; and second, MUs recruited at the same net neural drive in both contraction directions would not return two equal RT values if RT is quantified in absolute or relative force levels, as we described above. Because global EMG RMS reflects the summed synaptic drive to the whole MU pool rather than the input to any single unit, a given RT denotes the level of common pool drive at which that unit is recruited, not the intrinsic synaptic current its motoneuron requires to fire.

For each recorded trial consisting of six contractions, MU RTs were determined for each individual contraction ramp. The RT for a given MU and contraction was defined as the mean RMS value of the EMG signal within a 50 ms window centered on the MU’s first firing in that contraction. RT values with a *z*-score greater than 2 relative to the distribution of thresholds for that MU in that trial were discarded as outliers, as these likely reflected missed or incorrectly identified firings during manual editing, or large deviations in the angle of the performed contraction direction, that were observed in few contractions. Only MUs with identified and accepted recruitment thresholds in at least three of the six ramps were retained for further analysis. The individual contractiońs RTs for each MU were averaged and the resulting values sorted to obtain the recruitment order of all retained MUs for that trial.

The consistency of RTs and ROs were then compared between trials of the same contraction direction, as well as between contraction directions using Pearson correlation and statistical methods described in the Statistical Analysis section. We quantified the amount of relative RT change across trials by normalizing the absolute RT difference between the two trials with respect to trial 1, resulting in the normalized difference in recruitment threshold (ΔRT).

To quantify the number of MUs that exhibited a change in recruitment between contraction directions, we first quantified the test-retest variability of RT and RO between two trials of the same direction by computing the coefficient of repeatability according to Bland and Altman (1986). The coefficient of repeatability represents the 95% limits of agreement for repeated measurements. Because the RT and RO variability did not systematically depend on RT magnitude, we computed pooled coefficient of repeatabilitys (CRs) across the MUs of each participant. We then considered a MU to have changed its recruitment, if both its normalized ΔRT and change in RO position were above the respective participant-specific CRs values. This approach ensures that only changes greater than the expected measurement and physiological variability were interpreted as true alterations in recruitment behavior.

MU DRs were computed by averaging the median instantaneous discharge rate (IDR) during the steady plateau phase across all contractions, separately for abduction and flexion.

##### Motor Unit Coherence Analysis

To characterize the frequency-specific synchronization of common synaptic input to the MU pool in different contraction directions, intramuscular coherence (IMC) was computed on the motor unit data obtained from the coherence protocols. Only the steady-force plateau phases of the contractions were considered. As described above, MUs were tracked across contraction levels, resulting in concatenated files that include four contractions (two per contraction level). To correct for inconsistencies of the decomposition algorithm in detecting the exact discharge times of MUs, we realigned the MUs of each participant as described in Ibáñez et al. (2021). As the strength of common input is affected by the number of MU firings included in the cumulative spike trains (CSTs) (Farina et al. (2014)), which is in turn a consequence of the MU DRs given equal contraction durations, we ensured matching MU DRs across contraction directions by participant and direction-specific selection of contraction intensities (see methods above). We further included only MUs that were consistently firing throughout a minimum of two of the four contractions, and MUs with a z-score *<* 2 with respect to all active MUs of each file.

Coherence was computed on the CSTs of equally sized groups of MUs within each contraction direction. As increasing numbers of MUs per group results in higher coherence values, the number of MUs per group was half the total number of detected MUs in the contraction direction of less identified MUs to ensure comparability between directions. MUs were randomly assigned to the two groups with 100 repetitions per coherence computation. Coherence was calculated using MATLAB 2024b (MathWorks Inc.) using the function sp2a2 R2 mt within the Neurospec 2.11 toolbox (Halliday (2015)) in segments of length 1.6s and multi-tapers. We further transformed the coherence values into z-scores using the Fisher transform, scaled by the number of segments in the coherence computation (Laine et al. (2015)). For comparison of the z-coherence of the MU pools between the contraction direction, the area-under-curve (AuC) was computed for each participant in both contraction direction within the delta (1-5 Hz), alpha (5-13 Hz) and beta band (13-30 Hz). Only z-coherence values greater than the bias (mean z-score value between 250 and 500 Hz) were included in the AuC calculation.

### Statistical Analysis

We tested for a significant difference in MU recruitment threshold changes compared between trials of the same contraction direction and between abduction and flexion trials. We computed the median normalized absolute ΔRT value within MUs of each participant, once between trials of the same contraction direction (abduction - abduction, flexion - flexion) and between trials of different contraction directions (abduction - flexion). Normality of the paired differences for each participant’s resulting values was confirmed using Shapiro-Wilk test (p=0.72). Significance was then tested using a paired t-test. Effect sizes were expressed as Cohen’s d for paired samples, and 95% confidence intervals of the mean difference are reported.

To check for the relation between MU RT and DR and potential differences of such relation between contraction directions, *R*^2^ values of MU RT and DR were computed for each participant individually in both abduction and flexion. A Wilcoxon Signed-Rank Test was used to determine if the correlation between MU RT and DR (*R*^2^) differed significantly between conditions. To examine the relationship between the shift in RT and the adjustment in DR across contraction directions, we performed a one-way ANCOVA to determine the common within-subject association between ΔRT and difference in discharge rate (ΔDR), accounting for the dependence of observations within participants.

The difference in the strength of synchronization of synaptic input to FDI MUs between abduction and flexion was checked via the AuC of the z-coherence within the delta band (1-5), alpha band (5-13 Hz) and the beta band (13-30 Hz). Normality of the AuC values was confirmed using Shapiro-Wilk test (p=0.91) and significance checked using a paired t-test. Effect sizes were expressed as Cohen’s d for paired samples, and 95% confidence intervals of the mean difference are reported.

Where applicable, all statistical analyses were two-sided. A significance threshold of p *<* 0.05 was applied (* *<* 0.05, ** *<* 0.01, *** *<* 0.001). All statistical analyses were performed in MATLAB 2024b (MathWorks Inc.)

## Results

### Abduction and flexion activated the same FDI muscle territory

The spatial distribution of EMG amplitude across both the intramuscular array and the surface grid was qualitatively similar between abduction and flexion. Activation maps of a representative participant showed comparable amplitude topographies for the two contraction directions in both modalities, with no apparent shift of the active region between tasks (Fig. 1 C, D). The CoG shift between abduction and flexion did not significantly exceed the shift observed between two repetitions of the same contraction direction, for either the intramuscular array (within-direction: 0.25 *±* 0.24 mm; between-direction: 0.33 *±* 0.31 mm; Wilcoxon signed-rank, p = 0.88) or the surface grid (within-direction: 0.50 *±* 0.35 mm; between-direction: 0.64 ± 0.60 mm; p = 1.00; n = 6). The displacement of the activation center across contraction directions was therefore within the range of the trial-to-trial variability of the same contraction direction, indicating that abduction and flexion activated the same muscle territory rather than spatially distinct regions or muscle fibers.

### Motor Unit Decomposition

We implanted a 16-channel intramuscular thin-film electrode array in the FDI muscle, allowing us to decode and track the activity of larger numbers of MUs in a hand muscle compared to traditional intramuscular fine-wire or needle electromyography (EMG) or state-of-the-art high-density surface-EMG. For the recruitment protocol, we were able to identify per-participant averages of 23.3 *±* 5.5 MUs for index finger abduction and 23.0 *±* 7.0 MUs in flexion. Out of these MUs, 19.8 *±* 7.0 were identified in both contraction directions. In the coherence protocol, similar average MU numbers of 22.8 *±* 5.2 in abduction and 22.7 *±* 6.9 in flexion were decomposed, with 17.8 *±* 6.1 MUs tracked across both contraction directions. Tracked MUs showed a median *R*^2^ of 0.98 (IQR 0.95 - 0.99, n=120) against a null-distribution of all not-tracked MU pairs of median *R*^2^ = 0.75 (IQR 0.51 - 0.85, n=3343). As intended and ensured through the calibration process to determine participant-specific target force amplitudes for each contraction direction (see Methods), MU DRs were similar across abduction and flexion for each participant. In the recruitment protocol, average MU DR was 12.86 *±* 1.88 spikes/s during index finger abduction and 13.02 *±* 2.00 spikes/s in flexion. In the coherence protocol, values were similar with 13.76 *±* 1.91 spikes/s and 13.94 *±* 2.53 spikes/s in abduction and flexion respectively (Fig. 4B).

We did not find evidence of separate MU pools or single MUs recruited exclusively in one contraction direction. In few cases, we found clear absence of motor unit activity in one direction and clear presence of activity in the other. However, these absences likely reflect contraction direction-specific recruitment thresholds not being reached rather than true direction-specific activation.

### Reliable recruitment order within, but not across contraction directions

Figure 2A illustrates the process of RT extraction and the observed recruitment variability within and across contraction directions. Across all six participants, FDI MUs showed highly consistent RTs and consequently ROs between two trials of the same contraction direction. Figure 2B shows one exemplary participant’s ROs and RTs of two trials of abduction and flexion plotted on the x-axes (trial 1) and y-axes (trial 2) with the respective Pearson correlation coefficients. The narrow spread around the diagonal indicates minimal changes of RT within the same force direction. We observed MUs mostly kept their position within the RO, with rare deviations of 1-2 positions in the RO across the full dataset. Pearson correlation was high across all participants within the abduction (r_RT_ = 0.96 *±* 0.05, r_RO_ = 0.95 *±* 0.05) and flexion contractions (r_RT_ = 0.93 *±* 0.07, r_RO_ = 0.91 *±* 0.08). By contrast, recruitment between abduction and flexion revealed much less consistent RTs and ROs (Fig. 2C) (r_RT_ = 0.46 *±* 0.38, r_RO_ = 0.47 *±* 0.38). While some MUs still showed similar RT values in both contraction directions, others were recruited at drastically different amplitudes of net neural drive, consequently also dramatically altering their position in the RO. This behavior was observed in either direction of RT change, meaning some MUs were recruited earlier in abduction compared to flexion, while others showed the opposite. All participant-level plots of ROs and RTs across and within contraction direction can be found in supplementary figure S3. Figure 2A illustrates the reversal of RO between two exemplary MUs. During index finger abduction, MU#12 was consistently recruited after MU#9, while in flexion MU#9 was recruited after MU#12.

**Figure 2:**
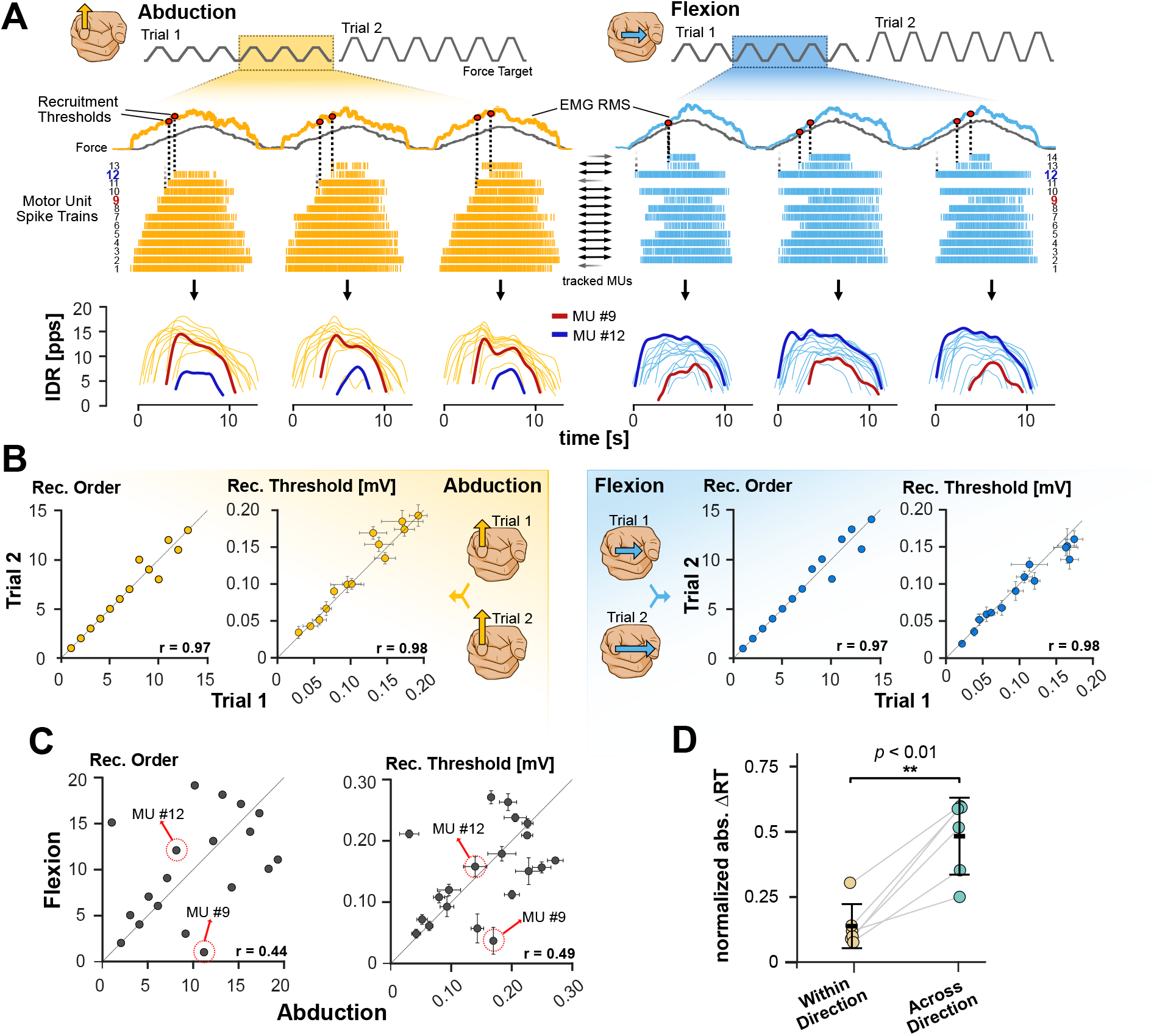
Recruitment variability within and across contraction directions. **A:** FDI MU recruitment thresholds and orders were computed via averaging the level of EMG activity at each MUs first spike across multiple repeated contractions separately for both contraction directions. Smoothed spike trains show a reversal of recruitment order of the exemplary MUs#9 and #12 between abduction and flexion. The change in recruitment is accompanied by a corresponding change of DR. **B:** MU recruitment thresholds and orders between two trials of abduction (left) and flexion (right). Errorbars indicate standard deviation across contractions in each trial. Pearson R indicates highly consistent RTs and ROs within the same contraction direction. **C:** RT and RO between abduction and flexion. Data shows some MUs considerably change their RTs and position in the RO between the two contraction directions, resulting in lower Pearson correlation values. MU#9 and MU#12 are highlighted relating to panel A. **D:** Each participant’s median absolute ΔRT within and across contraction directions. Lines connect same participant’s values. Errorbars show mean ± SD. ΔRT was significantly higher across than within directions (paired t-test, p = 0.0016). * p *<* 0.05, ** p *<* 0.01, *** p *<* 0.001

We quantified the relative change in recruitment threshold across contraction directions by computing the normalized absolute ΔRT. Across participants, absolute ΔRT values were significantly increased from within a contraction direction versus across contraction directions as quantified with the average of all participant’s median ΔRTs values (average median ΔRT within direction = 13.5 %, average median ΔRT across directions = 46.7 %. Paired t-test: t(5) = 6.16, p = 0.0016. Cohen’s d = 2.51, 95% CI [0.19, 0.47]) (Fig. 2D). The proportion of MUs that exhibited a clear change in recruitment was determined by employing the coefficient of repeatability of the normalized ΔRT and the change in RO position as criteria for recruitment change (see Methods). Within the same functional task (i.e., identical contraction direction), only 5.0% of MUs across all participants showed a change in recruitment, whereas 45.3% changed their recruitment across contraction directions.

### Coordinated Modulation of Recruitment and Discharge Rate

In accordance with the onion-skin principle, we consistently found the MU DRs to be inversely correlated to the specific MUs RT across both contraction directions (Fig. 3A, B), meaning MUs with higher RTs had lower steady-state DRs than lower-threshold MUs. Importantly, the strength of RT-DR correlation did not depend on contraction direction, as we did not find any significantly higher correlation of RT and DR in either contraction direction with average *R*^2^ values of 0.44 *±* 0.18 in abduction and 0.48 *±* 0.10 in flexion at the lower force level, and 0.22 *±* 0.10 and 0.24 *±* 0.24 at the higher force level (Wilcoxon signed-rank test, F1: Z=-0.94, p=0.35, F2: Z=-0.31, p=0.75). Because the global EMG and consequently the mean DR were matched across tasks, the overall drive to the pool was comparable in abduction and flexion. A shift in an individual unit’s RT therefore reflects a change in the share of this common drive that the unit receives, i.e. a redistribution of synaptic input across the pool, rather than a change in its intrinsic recruitment threshold. In line with this, the change in each MU’s RT was mirrored by an opposite change in its steady-state DR (Fig. 3C,D). MUs recruited at lower pool drive in flexion also discharged faster, and vice versa, as expected when a motoneuron receives more or less synaptic input. This co-modulation preserved the RT–DR relation despite the large RT changes.

**Figure 3:**
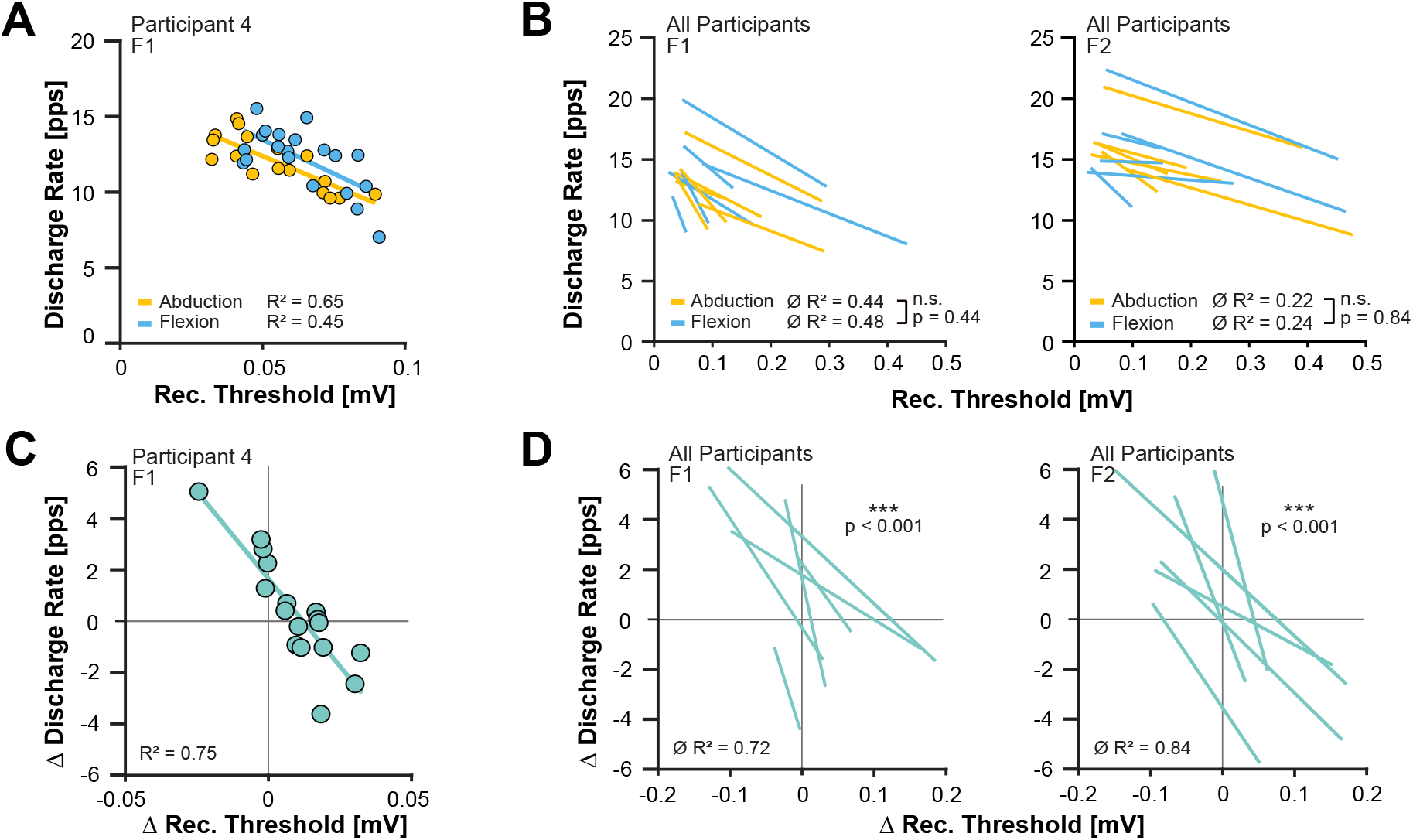
Modulation of DR and RT across contraction directions. **A:** Exemplary participant’s data, showing inverse correlation of DR and RT with linear regression lines. **B:** Correlation between RT and DR split into lower force (left, “F1”) and higher force (right, “F2”) trials showing all participants linear regression lines. No significant different correlation was found between abduction and flexion (Wilcoxon Signed-Rank test, F1 p = 0.35, F2 p = 0.75). **C:** Same MUs as in A. Difference in DR over difference in RT between abduction and flexion with corresponding linear regression. MUs that reduce their RT show increased DR and vice versa. **D:** Difference in DR over difference in RT split into lower and higher force trials showing all participants linear regression lines. A highly significant correlation of differences in RT and DR is found across participant in both force levels (repeated-measures correlation, p*<*0.001). * p *<* 0.05, ** p *<* 0.01, *** p *<* 0.001.

This behavior was observed across all participants and both force levels (Fig. 3C, D) with strong statistical significance (mean *R*^2^ F1= 0.72 *±* 0.14, mean *R*^2^ F2= 0.84 *±* 0.11, repeated-measures correlation, F1: p*<*0.001, 95% CI [-4.86, 6.93], F2: p*<*0.001, 95% CI [-3.91, 5.96]). All participant level plots analogue to figure 3A, C can be found in supplementary figure S4.

### FDI motor units show stronger beta band coherence in abduction than flexion

We characterized the synchronization of common synaptic input to the motoneuron pool of the FDI muscle during index finger abduction and flexion by calculating IMC. For each participant, FDI MUs were randomly divided into two subgroups and combined into CSTs before coherence analysis, with 100 random permutations of group assignment (Fig. 4A).

**Figure 4:**
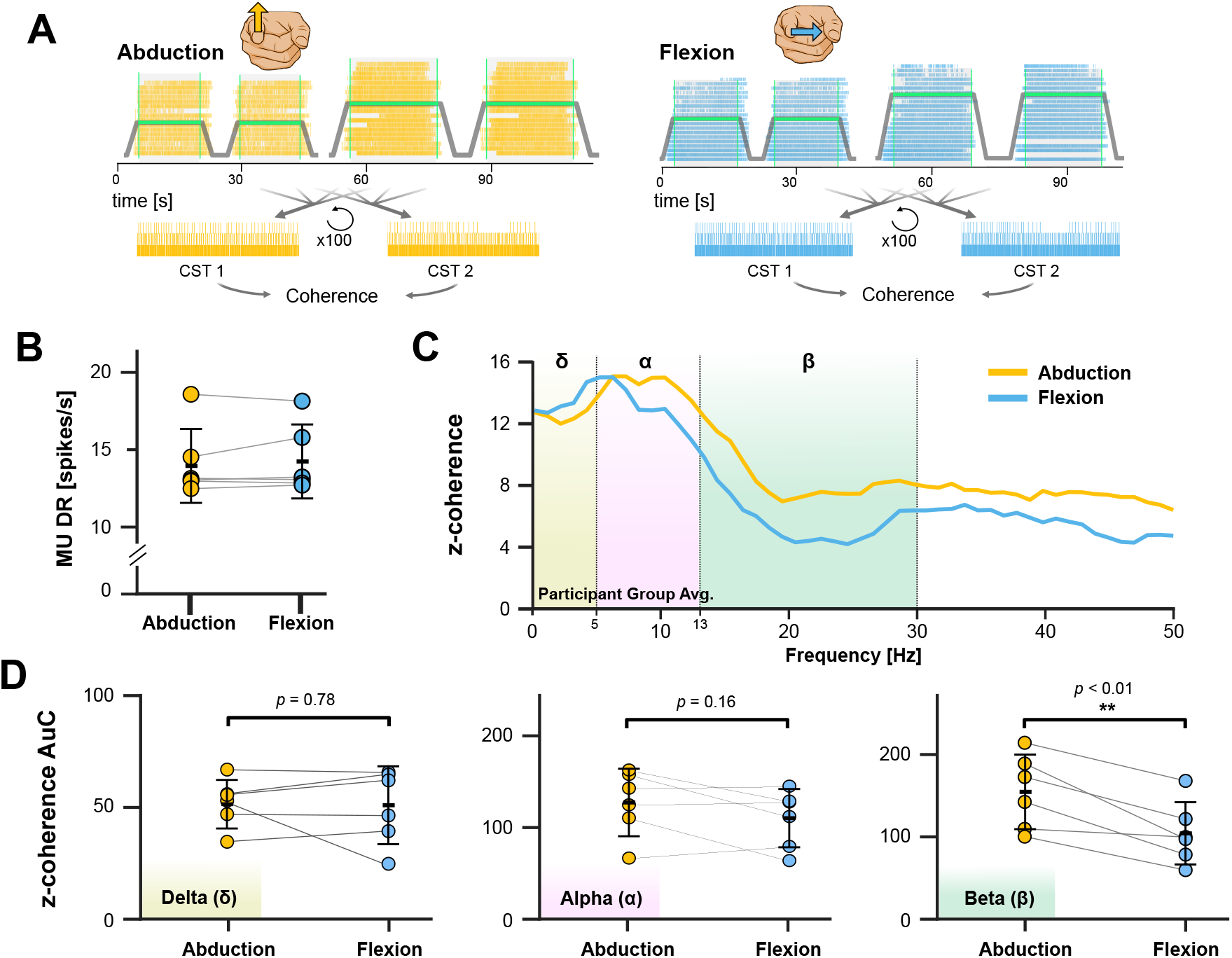
Intramuscular coherence in abduction compared to flexion. **A:** Coherence is calculated between two randomly selected subgroups of FDI MUs per contraction direction and averaged across 100 permutations, considering only firings during steady plateau phases. **B:** MU DRs in coherence protocol data across participants. **C:** Participant group average intramuscular coherence. Intramuscular z-coherence in abduction and flexion with delta (1-5 Hz), alpha (5-13 Hz) and beta (13-30 Hz) bands highlighted. MUs show stronger beta band coherence during abduction. **D:** Across-participants coherence in delta (left), alpha (center) and beta (right) bands, expressed as area-under-curve above bias. No significant difference was observed in delta and alpha, whereas beta-band coherence was significantly reduced in flexion compared to abduction. * p *<* 0.05, ** p *<* 0.01, *** p *<* 0.001.

To avoid confounding effects of differing motor unit DRs across contraction directions, we employed a dedicated calibration protocol to select participant- and direction-specific target force levels that matched global EMG activity between abduction and flexion (see Methods). As a result, MU firing rates were highly similar across directions in all participants (Fig. 4B). This step was crucial, since mismatched firing rates at identical contraction durations would otherwise result in unequal numbers of MU discharges contributing to the coherence estimate, potentially biasing the results (Negro and Farina (2012); Negro et al. (2009); Farina et al. (2014); De la Rocha et al. (2007)).

The strength of common input synchronization was then compared between contraction directions using z-transformed coherence values (z-coherence). We analyzed IMC separately within the delta (1-5 Hz), alpha (5-13 Hz), and beta band (13-30 Hz). Figure 4C shows the intramuscular z-coherence during index finger abduction and flexion averaged across all participants, with noticeably greater coherence values in the beta band from 13-30 Hz during abduction. Across participants, beta-band z-coherence was significantly lower during flexion than abduction (t(5) = 4.59, p = 0.006, Cohen’s d = 1.87, 95% CI [21.79, 77.24]). By contrast, no significant difference in coherence between contraction directions was found in the delta band (Fig. 4D, t(5) = 0.30, p = 0.776, Cohen’s d = 0.12, 95% CI [-10.70, 13.54]) and alpha band (t(5) = 1.64, p = 0.162, Cohen’s d = 0.56, 95% CI [-9.85, 44.47]).

## Discussion

The primary finding of this study is the significant flexibility in the distribution of neural drive to the MUs of the FDI muscle depending on the functional task. In the MU activity extracted from HD-iEMG in six healthy participants, we found that 45.3% of MUs substantially changed their recruitment thresholds (RT) and recruitment order (RO) when switching between abduction and flexion (RT and RO above the participant’s coefficient of repeatability). This variability in recruitment was far above natural variability in MU discharge timings or recruitment thresholds across multiple contractions, as shown by the highly consistent RTs and ROs that we observed between contractions in the same contraction direction with only 5.0% of MUs above the CRs (average ΔRT across directions = 46.7%, within directions = 13.5 %) (Figs. 2B,C,D). These findings align with previous observations in the FDI (Desmedt and Godaux (1981); Thomas et al. (1986)) and demonstrate that MU recruitment cannot be fully explained by a rigid application of Henneman’s size principle to the entire anatomical MU pool across different functional tasks. Instead, the CNS appears capable of adjusting the distribution of neural drive to subsets of MUs within the pool, likely through specific excitatory or inhibitory synaptic currents. This does not contradict the size principle itself, which constrains recruitment among motoneurons receiving a common input rather than fixing one ordering across an entire anatomical pool (Bawa et al. (2014); Hug et al. (2023)). Under this view, the reordering we observe implies that the FDI MUs we sampled do not all receive the same input in both tasks, with the relative weighting of common inputs to subsets of the pool changing between abduction and flexion. Note that Desmedt and Godaux (1981) expressed recruitment thresholds in absolute force, whereas we quantify them in global EMG amplitude. Since FDI EMG at a matched absolute force is lower in flexion than in abduction (Supplementary Fig. S2B), thresholds expressed in force would systematically overstate the apparent threshold increase from abduction to flexion, which is why we report thresholds as a measure of the drive to the muscle.

Despite the observed variability in RTs, the fundamental control properties of the identified MU pool remained stable. In both index finger abduction and flexion, we found statistically similar negative correlations between RT and DR, where later recruited MUs exhibited lower DRs than those recruited earlier (Fig. 3A,B). This hierarchical organization has been described as the onion-skin scheme and has been reported in various conditions at low-to-medium force levels (De Luca et al. (1982); De Luca and Hostage (2010); Hu et al. (2014); Tanji and Kato (1973)). The RT-DR relation was weaker at the higher force level (Fig. 3B), which is in agreement with previous studies and may be caused by the saturation of DRs of low-threshold MUs (De Luca and Hostage (2010); Hu et al. (2014)).

Notably, MUs that shifted their RT from abduction to flexion showed a corresponding opposite modulation of their steady-state DR (Fig. 3C,D), preserving the RT-DR relation even though large changes in RT were present. Because comparable global EMG indicates similar total drive to the pool across tasks, these RT shifts are most likely explained by a redistribution of common synaptic input, with individual units receiving a larger or smaller share depending on the task, rather than by changes in the intrinsic recruitment thresholds of the motoneurons. That the RT-DR relation is preserved through this redistribution indicates the onion-skin organization of MU firing reflects common synaptic input to the pool rather than intrinsic motoneuron properties, while recruitment order itself remains task-dependent.

The HD-iEMG electrode arrays used in this study enable the concurrent sampling of large numbers of MUs, allowing for correlation-based measures in the frequency domain (Negro and Farina (2012)), specifically intramuscular coherence to characterize the frequency-specific synchronization of common input to the FDI MUs across the two tasks. Coherence reflects the synchronization of shared input rather than its net magnitude and does not by itself identify the source of that input. The magnitude of net neural drive is instead addressed separately, through the amplitude matching and the recruitment-threshold and discharge-rate data. We hypothesized that the FDI might receive more strongly synchronized common input when it acts as a prime mover (abduction) than when it functions as a synergist during index finger flexion, where a greater involvement of spinal circuits shared with the long finger flexors could reduce the synchronization of high-frequency input. This could be facilitated by spinal premotor interneuronal circuits (He et al. (2025)). Within the delta band, which corresponds to low-frequency correlation from 1-5 Hz and reflects the slow, common synaptic input that drives force fluctuations and is directly related to muscle force production and function (Farmer et al. (1993); De Luca and Erim (1994); Negro et al. (2009)), no significant difference in strength of coherence was found. Similarly, the alpha band between 5 - 13 Hz, related to sensory afferents, muscle spindle feedback coupling and includes the physiological tremor frequency (Christakos et al. (2006); Farina et al. (2017); Laine and Valero-Cuevas (2017)), did not show significant difference.

We did find a significantly lower FDI MU coherence during index finger flexion compared to abduction in the beta band from 13 - 30 Hz (Fig. 4C,D). Because this difference was confined to the beta band, it is unlikely to arise from a global effect of drive magnitude, MU count, or discharge rate, which would be expected to shift coherence across frequencies rather than selectively in the band associated with corticospinal pathways (Conway et al. (1995); Salenius et al. (1997); Baker et al. (1999); Ibáñez et al. (2021)). We therefore interpret the reduced beta-band synchronization during flexion as consistent with a reduced or less synchronized corticospinal contribution when the FDI acts as a synergist, possibly reflecting a greater involvement of spinal circuits shared with the long finger flexors. We emphasize, however, that this is a hypothesis rather than a demonstrated mechanism as intramuscular coherence reflects the synchronization of shared input, not its magnitude, and cannot identify the source of that input. The observed difference could also reflect other factors, such as increased synaptic noise or changes in the synchronization of the MU population. Our design establishes that FDI motor unit recruitment reorganizes between abduction and flexion, but it cannot isolate the cause of this reorganization. The two tasks differ not only in the functional role of the FDI (prime mover vs. synergist) but also in their mechanical demands and in the involvement of the long finger flexors as additional prime movers, and these factors cannot be disentangled here. Resolving which of them drives the reorganization, and clarifying its neural determinants, in particular the reduced beta-band synchronization, will require simultaneous recordings from FDI and the finger flexors (FDP and FDS) during matched tasks, ideally combined with concurrent EMG and EEG to assess corticomuscular coherence

We found no evidence of distinct “task-specific” motor unit pools in the FDI, which is consistent with previous reports (Thomas et al. (1986)). A potential confound of matching global EMG amplitude across tasks is that a comparable amplitude could, in principle, arise from spatially distinct regions of the FDI, and thus from different MU populations being engaged in abduction and flexion, innervating different sets of muscle fibers. To address this, we quantified the spatial distribution of muscle activation as the amplitude-weighted CoG of the RMS activation maps, both within the intramuscular array and across the high-density surface-EMG (HD-sEMG) grid. In both modalities, the displacement of the CoG between abduction and flexion did not significantly exceed the displacement observed between two repetitions of the same contraction direction (Fig. 1 D), indicating that the two tasks activated the same muscle territory rather than spatially separate regions. Accordingly, in the few instances where a MU was identified in only one contraction direction, it is most likely that its direction-specific recruitment threshold was simply not reached in the respective other direction, given the large changes in RT we observed between directions (mean RT change across contraction directions of 46.7%), rather than a distinct population being engaged.

These findings are in line with the existing literature on regional FDI activation. Earlier work reporting task- and direction-dependent differences in FDI activity describes a reduction in surface EMG amplitude toward index finger flexion (Whitford and Kukulka (2011); Zhou et al. (2011)), and direction-dependent MU twitch and force vectors (Suresh et al. (2012)), but does not demonstrate a shift in the center of activation, the engagement of distinct muscle regions, or the recruitment of a separate MU population across contraction directions. Together with our own spatial analysis, this supports controlling for EMG amplitude across contraction directions as a valid proxy for comparable net neural drive, and indicates that the limited spatial sampling of the intramuscular array is not a confounding factor for the observed recruitment variability.

Our experimental setup utilized a 2D dynamometer with real-time feedback to strictly control contraction vectors, eliminating voluntary or involuntary deviation in the applied force vector as a confounding factor (Supplementary Fig. S1B). However, certain limitations exist, such as that the contractions were performed at relatively low forces (10-20% MVC). While this range captures a representative distribution of the FDI motor pool, where the upper limit of recruitment range is reported to be even below 50% maximum abduction force, with most MUs even recruited below 30% (Duchateau and Enoka (2022); De Luca et al. (1982)), we cannot extrapolate these findings to the highest-force conditions.

In summary, the present study provides further insights into the distribution of net neural drive within a pool of MUs across different functional tasks. Using current state-of-the-art HD-iEMG technology, we observed significant changes in recruitment thresholds in the FDI muscle in index finger abduction and flexion with corresponding adaptations in rate coding. We further quantified the intramuscular coherence within the observed pool of MUs and found significantly decreased beta coherence during index finger flexion compared to abduction, indicating reduced synchronization of common input in this band when the muscle acts as a synergist. This work further provides the basis for future investigations in the role of underlying corticospinal mechanisms and their alterations in disease.

## Acknowledgements

M.O. acknowledges the Deutsche Forschungsgemeinschaft (DFG, German Research Foundation) - Project 523352235. J.I. acknowledges the European Research Council (ERC) - Project ECHOES - 0107769. M.R. acknowledges the German Federal Ministry of Research, Technology and Space (ACS iIMMUNE, 01EO2105) and Deutsche Forschungsgemeinschaft (DFG, German Research Foundation) - Project SFB 1483-442419336

---

**Figure S1.**
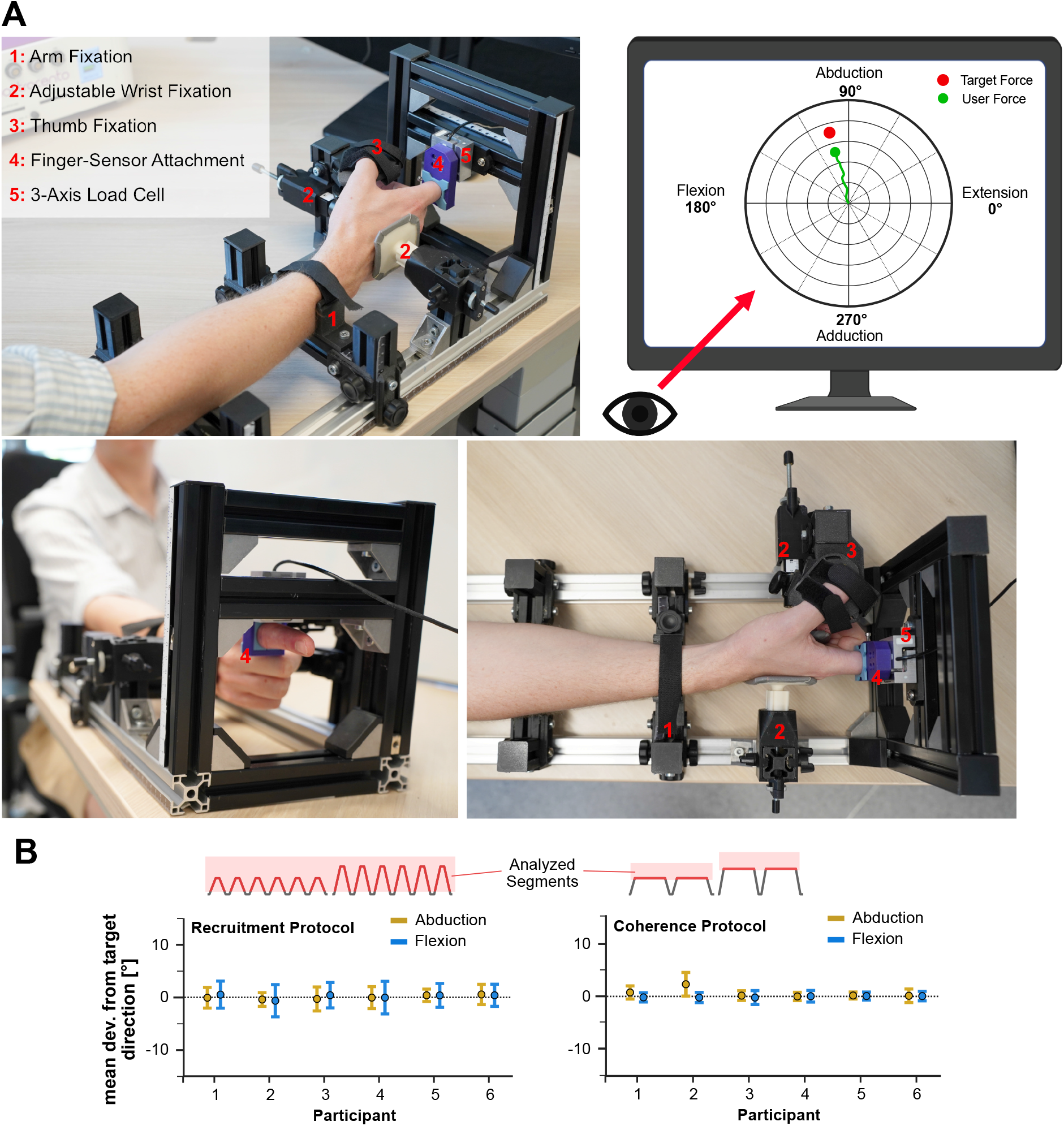
Detailed Experimental Setup. A: Custom Dynamometer for 2D isometric index finger contractions. Main features marked with red numbers and described. The setup features a 3-axis load cell capable of sensing the applied index finger force in all possible degrees of freedom. The hand and forearm positions are fixed with adjustable restraints, thereby fixing the wrist position and restricting force application through wrist contractions. The thumb position is likewise fixed to ensure a constant posture throughout an experimental protocol. Top right: 2D force feedback as observed by the user, displaying the user’s force as a green dot and trajectory with force amplitude as distance from center and force direction as angular position. Target force is displayed in red. B: Analysis of all participant’s directional accuracy throughout the contractions of the experimental protocols of the present study. Mean and standard deviation of the deviation of applied force to target force is shown, split into abduction and flexion contractions. For the recruitment protocol (left) the applied force directions of the full ramp contractions, including the concentric phase is considered. For the coherence protocol, only the force applied during the plateau phase is considered. In both protocols, force application was precise across participants, showing mean deviations close to 0 with low standard deviations. Mean deviations range from -0.85° to 0.51° in the recruitment protocol and -0.21 to 2.30° in the coherence protocol.

**Figure S2.**
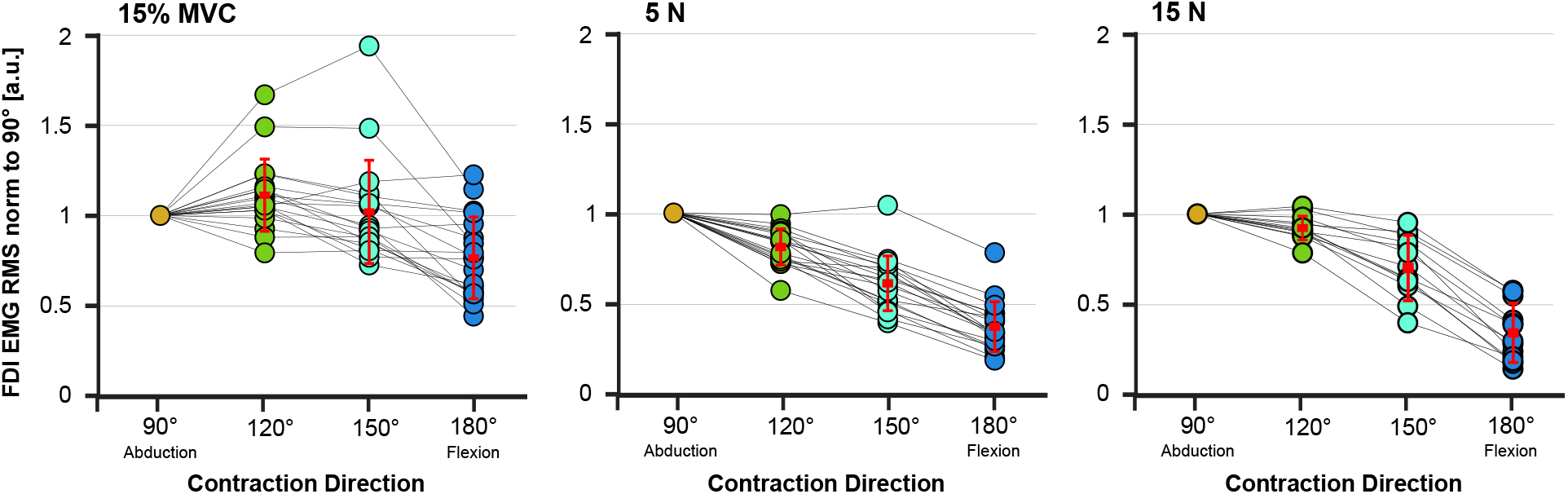
FDI activity amplitude in different contraction directions relative to abduction. Across participants, the plots show FDI muscle activity levels relative to the abduction activity during different contraction directions at matched relative force levels (left) and at identical absolute force levels at 5N and 15 N (center and right) per direction. In neither case did FDI activity levels match across directions, indicating the need for participant- and direction-specific target force amplitudes for the present study protocol. Dots represent each participant’s FDI EMG amplitudes with lines connecting the values corresponding to single individuals across force directions. **Methods:** 19 healthy participants (age 26.8 *±* 4.5 years. 14 male, 5 female) performed isometric ramp contractions in index finger abduction (90°), flexion (180°) and two directions equally spaced in between (120° and 150°). Participants used the dynamometer setup described in the main text and Fig. S1. Maximum voluntary contractions (MVC) values were acquired for each direction separately. Participants performed ramp contractions at 15% of the respective MVC in each direction (Protocol 1), as well as contractions at consistently increasing force until failure (Protocol 2). High-Density surface EMG signals (GR03MM1807, OT Bioelettronica, Turin, Italy) were captured from the FDI muscle. EMG amplitude was calculated as the mean root-mean-square (RMS) value across the full EMG grid during the contractiońs plateau phase for the 15% MVC ramp contractions (Protocol 1, left plot), and within a 10% force window around the 5N and 15N absolute forces (Protocol 2, center and right plot). All values were then normalized to the respective participant’s value in index finger abduction.

**Figure S3.**
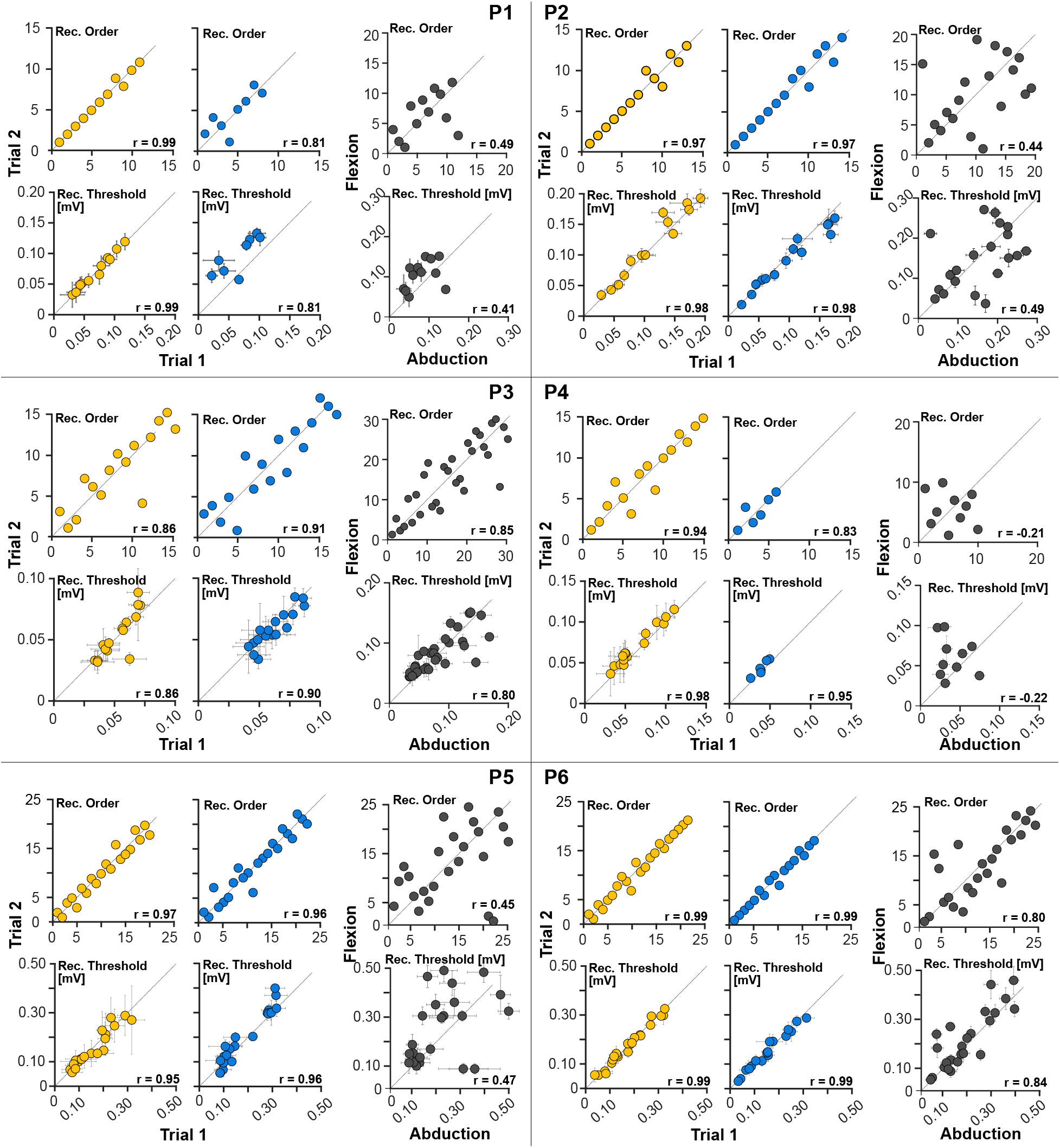
Participant-level recruitment variability within and across contraction directions. For each participant, plots show the recruitment thresholds in mV, and recruitment orders between two trials of the same contraction direction (Yellow: Abduction, blue: Flexion) and between abduction and flexion (right column).

**Figure S4.**
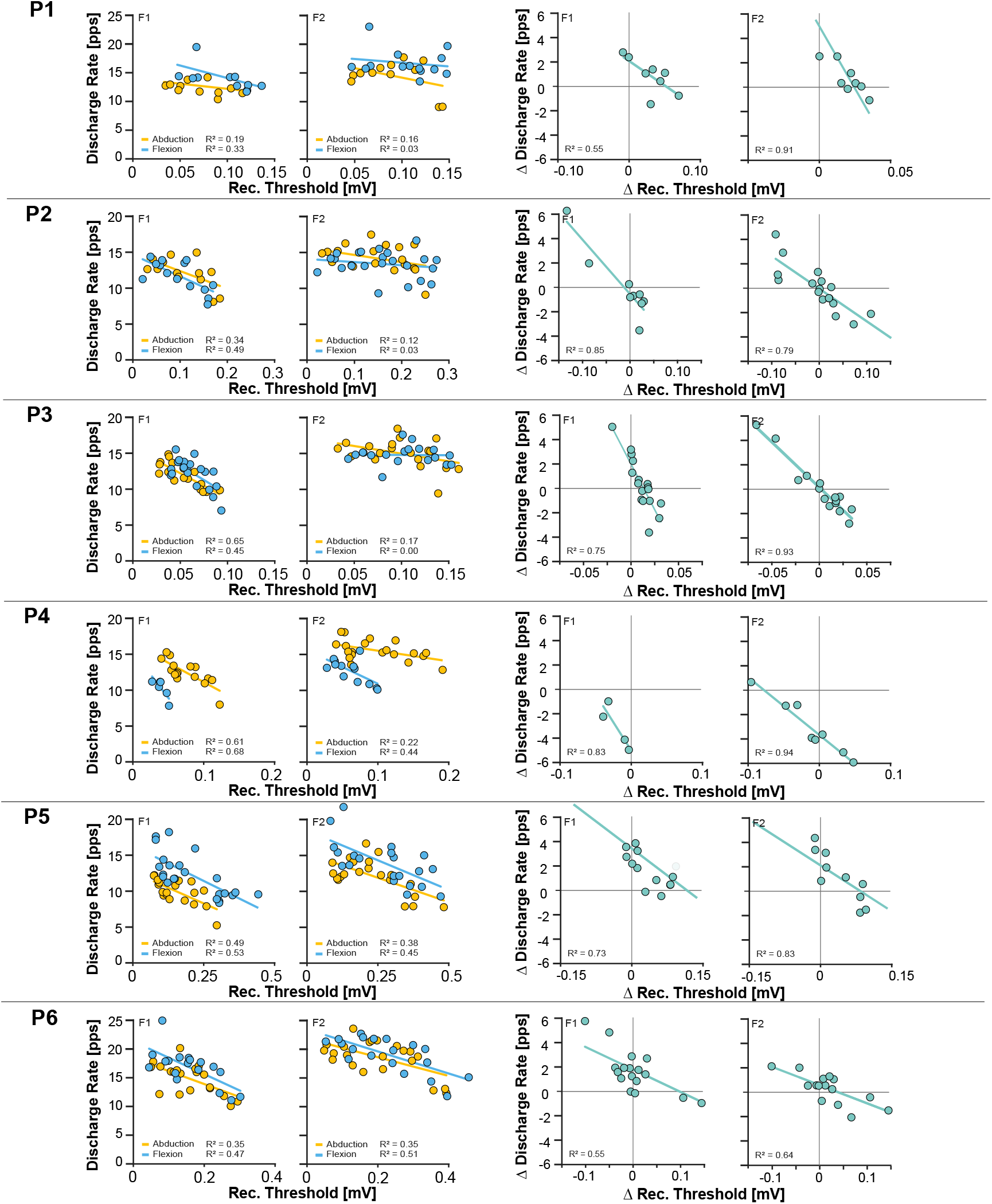
Participant-level modulation of discharge rate and recruitment threshold within and across contraction directions. For each participant, plots show the discharge rate in pps against the recruitment thresholds in mV separately for contraction directions and contraction levels (left), as well as the difference in discharge rate over the difference in recruitment threshold from abduction to flexion, separately for the two contraction levels (right). Lines correspond to the linear regression per dataset.

## References

Avrillon S, Hug F, Baker SN, Gibbs C, Farina D (2024a) Tutorial on MUedit: An open-source software for identifying and analysing the discharge timing of motor units from electromyographic signals. Journal of Electromyography and Kinesiology 77:102886.

Avrillon S, Hug F, Enoka RM, Caillet AH, Farina D (2024b) The identification of extensive samples of motor units in human muscles reveals diverse effects of neuromodulatory inputs on the rate coding. eLife 13:RP97085.

Baker SN, Kilner JM, Pinches EM, Lemon RN (1999) The role of synchrony and oscillations in the motor output. Experimental Brain Research 128:109–117.

Bawa PNS, Jones KE, Stein RB (2014) Assessment of size ordered recruitment. Frontiers in Human Neuroscience 8.

Bland J, Altman D (1986) Statistical methods for assessing agreement between two methods of clinical measurement. The Lancet 327:307–310.

Butler TJ, Kilbreath SL, Gorman RB, Gandevia SC (2005) Selective recruitment of single motor units in human flexor digitorum superficialis muscle during flexion of individual fingers. The Journal of Physiology 567:301.

Calancie B, Bawa P (1985) Voluntary and reflexive recruitment of flexor carpi radialis motor units in humans. Journal of Neurophysiology 53:1194–1200.

Campbell E, Egle F, Oßwald M, Cŏté-Allard U, Pilarski PM, Boccardo N, Meattini R, Vujaklija I, Hargrove L, Canepa M, Eddy E, Vecchio AD, Castellini C, Scheme E (2025) (Un)supervised (Co)adaptation via Incremental Learning for Myoelectric Control: Motivation, Review, and Future Directions. IEEE Transactions on Neural Systems and Rehabilitation Engineering 33:3565–3582.

Capsi-Morales P, Barsakcioglu DY, Catalano MG, Grioli G, Bicchi A, Farina D (2025) Merging motoneuron and postural synergies in prosthetic hand design for natural bionic interfacing. Science Robotics 10:eado9509.

Chen C, Yu Y, Sheng X, Farina D, Zhu X (2021) Simultaneous and proportional control of wrist and hand movements by decoding motor unit discharges in real time. Journal of Neural Engineering 18:056010.

Christakos CN, Papadimitriou NA, Erimaki S (2006) Parallel Neuronal Mechanisms Underlying Physiological Force Tremor in Steady Muscle Contractions of Humans. Journal of Neurophysiology 95:53–66.

Conway BA, Halliday DM, Farmer SF, Shahani U, Maas P, Weir AI, Rosenberg JR (1995) Synchronization between motor cortex and spinal motoneuronal pool during the performance of a maintained motor task in man. The Journal of Physiology 489:917–924.

Conwit RA, Stashuk D, Tracy B, McHugh M, Brown WF, Metter EJ (1999) The relationship of motor unit size, firing rate and force. Clinical Neurophysiology 110:1270–1275.

De la Rocha J, Doiron B, Shea-Brown E, Josić K, Reyes A (2007) Correlation between neural spike trains increases with firing rate. Nature 448:802–806.

De Luca CJ, LeFever RS, McCue MP, Xenakis AP (1982) Behaviour of human motor units in different muscles during linearly varying contractions. The Journal of Physiology 329:113–128.

De Luca CJ, Erim Z (1994) Common drive of motor units in regulation of muscle force. Trends in Neurosciences 17:299–305.

De Luca CJ, Hostage EC (2010) Relationship Between Firing Rate and Recruitment Threshold of Motoneurons in Voluntary Isometric Contractions. Journal of Neurophysiology 104:1034–1046.

Del Vecchio A, Holobar A, Falla D, Felici F, Enoka R, Farina D (2020) Tutorial: Analysis of motor unit discharge characteristics from high-density surface EMG signals. Journal of Electromyography and Ki-nesiology 53:102426.

Desmedt JE, Godaux E (1977) Fast motor units are not preferentially activated in rapid voluntary contractions in man. Nature 267.

Desmedt JE, Godaux E (1981) Spinal Motoneuron Recruitment in Man: Rank Deordering with Direction But Not with Speed of Voluntary Movement. Science 214:933–936.

Duchateau J, Enoka RM (2022) Distribution of motor unit properties across human muscles. Journal of Applied Physiology 132:1–13.

Farina D, Negro F, Dideriksen JL (2014) The effective neural drive to muscles is the common synaptic input to motor neurons. The Journal of Physiology 592:3427–3441.

Farina D, Vujaklija I, Sartori M, Kapelner T, Negro F, Jiang N, Bergmeister K, Andalib A, Principe J, Aszmann OC (2017) Man/machine interface based on the discharge timings of spinal motor neurons after targeted muscle reinnervation. Nature Biomedical Engineering 1:0025.

Farmer SF, Bremner FD, Halliday DM, Rosenberg JR, Stephens JA (1993) The frequency content of common synaptic inputs to motoneurones studied during voluntary isometric contraction in man. The Journal of Physiology 470:127–155.

Feiereisen P, Duchateau J, Hainaut K (1997) Motor unit recruitment order during voluntary and electrically induced contractions in the tibialis anterior. Experimental Brain Research 114:117–123.

Frančič A, Holobar A (2021) On the Reuse of Motor Unit Filters in High Density Surface Electromyograms Recorded at Different Contraction Levels. IEEE Access 9:115227–115236.

Garland SJ, Cooke JD, Miller KJ, Ohtsuki T, Ivanova T (1996) Motor unit activity during human single joint movements. Journal of Neurophysiology 76:1982–1990.

Grison A, Gibbs C, Rawji V, Vila LG, Szczech I, Varghese R, Bryan P, Kundu A, Yang X, Gallego J Farina D (2025) Multidimensional motoneuron control using intramuscular microelectrode arrays in tetraplegic spinal cord injury. medRxiv preprint medRxiv 2025.07.17.25331429.

Grison A, Pereda JI, Muceli S, Kundu A, Baracat F, Indiveri G, Donati E, Farina D (2024) Intramuscular High-Density Micro-Electrode Arrays Enable High-Precision Decoding and Mapping of Spinal Motor Neurons to Reveal Hand Control. arXiv preprint arXiv:2410.11016.

Halliday DM (2015) Nonparametric directionality measures for time series and point process data. Journal of Integrative Neuroscience 14:253–277.

He B, Salmas P, Zhang J, Duan X, Cheung VCK (2025) State-dependent neural representations of muscle synergies in the spinal cord revealed by optogenetic stimulation. The Journal of Physiology 603:4659–4679.

Henneman E, Clamann HP, Gillies JD, Skinner RD (1974) Rank order of motoneurons within a pool: law of combination. Journal of Neurophysiology 37:1338–1349.

Henneman E (1957) Relation between size of neurons and their susceptibility to discharge. Science (New York, N.Y.) 126:1345–1347.

Henneman E, Somjen G, Carpenter DO (1965) Functional significance of cell size in spinal motoneurons. Journal of Neurophysiology 28:560–580.

Holobar A, Zazula D (2007) Multichannel Blind Source Separation Using Convolution Kernel Compensation. IEEE Transactions on Signal Processing 55:4487–4496.

Hu X, Rymer WZ, Suresh NL (2014) Control of motor unit firing during step-like increases in voluntary force. Frontiers in Human Neuroscience 8.

Hudson AL, Taylor JL, Gandevia SC, Butler JE (2009) Coupling between mechanical and neural behaviour in the human first dorsal interosseous muscle. The Journal of Physiology 587:917–925.

Hug F, Avrillon S, Ibáñez J, Farina D (2023) Common synaptic input, synergies and size principle: Control of spinal motor neurons for movement generation. The Journal of Physiology 601:11–20.

Ibáñez J, Vecchio AD, Rothwell JC, Baker SN, Farina D (2021) Only the Fastest Corticospinal Fibers Contribute to β Corticomuscular Coherence. Journal of Neuroscience 41.

Laine CM, Martinez-Valdes E, Falla D, Mayer F, Farina D (2015) Motor Neuron Pools of Synergistic Thigh Muscles Share Most of Their Synaptic Input. The Journal of Neuroscience 35:12207–12216.

Laine CM, Valero-Cuevas FJ (2017) Intermuscular coherence reflects functional coordination. Journal of Neurophysiology 118:1775–1783.

Liss FE (2012) The Interosseous Muscles: The Foundation of Hand Function. Hand Clinics 28:9–12.

Maillet J, Avrillon S, Nordez A, Rossi J, Hug F (2022) Handedness is associated with less common input to spinal motor neurons innervating different hand muscles. Journal of Neurophysiology 128:778–789.

Marshall NJ, Glaser JI, Trautmann EM, Amematsro EA, Perkins SM, Shadlen MN, Abbott LF, Cunningham JP, Churchland MM (2022) Flexible neural control of motor units. Nature Neuroscience 2022 25:11 25:1492–1504.

Masakado Y, Akaboshi K, Nagata Ma, Kimura A, Chino N (1995) Motor unit firing behavior in slow and fast contractions of the first dorsal interosseous muscle of healthy men. Electroencephalography and Clinical Neurophysiology/Electromyography and Motor Control 97:290–295.

Masquelet AC, Salama J, Outrequin G, Serrault M, Chevrel JP (1986) Morphology and functional anatomy of the first dorsal interosseous muscle of the hand. Surgical and Radiologic Anatomy 8:19–28.

Milner-Brown HS, Stein RB, Yemm R (1973) The orderly recruitment of human motor units during voluntary isometric contractions. The Journal of Physiology 230:359–370.

Muceli S, Poppendieck W, Negro F, Yoshida K, Hoffmann KP, Butler JE, Gandevia SC, Farina D (2015) Accurate and representative decoding of the neural drive to muscles in humans with multi-channel intra-muscular thin-film electrodes. The Journal of Physiology 593:3789–3804.

Negro F, Farina D (2012) Factors Influencing the Estimates of Correlation between Motor Unit Activities in Humans. PLoS ONE 7:e44894.

Negro F, Holobar A, Farina D (2009) Fluctuations in isometric muscle force can be described by one linear projection of low-frequency components of motor unit discharge rates. The Journal of physiology 587:5925–5938.

Negro F, Muceli S, Castronovo AM, Holobar A, Farina D (2016) Multi-channel intramuscular and surface EMG decomposition by convolutive blind source separation. Journal of neural engineering 13.

Oßwald M, Cakici AL, Souza de Oliveira D, Braun DI, Farina D, Del Vecchio A (2025) Task-specific motor units in the extrinsic hand muscles control single- and multidigit tasks of the human hand. Journal of Applied Physiology 138:1187–1200.

Ranney D, Wells R (1988) Lumbrical muscle function as revealed by a new and physiological approach. The Anatomical Record 222:110–114.

Riek S, Bawa P (1992) Recruitment of motor units in human forearm extensors. Journal of Neurophysiology 68:100–108.

Saboisky JP, Butler JE, Fogel RB, Taylor JL, Trinder JA, White DP, Gandevia SC (2006) Tonic and Phasic Respiratory Drives to Human Genioglossus Motoneurons During Breathing. Journal of Neurophysiology 95:2213–2221.

Salenius S, Portin K, Kajola M, Salmelin R, Hari R (1997) Cortical Control of Human Motoneuron Firing During Isometric Contraction. Journal of Neurophysiology 77:3401–3405.

Scutter SD, Türker KS (1998) Recruitment stability in masseter motor units during isometric voluntary contractions. Muscle & Nerve 21:1290–1298.

Stein RB, Gossen ER, Jones KE (2005) Neuronal variability: noise or part of the signal? Nature Reviews Neuroscience 6:389–397.

Stotz PJ, Bawa P (2001) Motor unit recruitment during lengthening contractions of human wrist flexors. Muscle & Nerve 24:1535–1541.

Suresh N, Kuo A, Heckman C, Rymer WZ (2012) Using spike-triggered averaging to characterize motor unit twitch vectors in the first dorsal interosseous In 2012 Annual International Conference of the IEEE Engineering in Medicine and Biology Society, pp. 3604–3607 ISSN: 1558-4615.

Tanji J, Kato M (1973) Firing rate of individual motor units in voluntary contraction of abductor digiti minimi muscle in man. Experimental Neurology 40:771–783.

Tax AAM, van der Gon JJD, Gielen CCAM, van den Tempel CMM (1989) Differences in the activation of m. biceps brachii in the control of slow isotonic movements and isometric contractions. Experimental Brain Research 76:55–63.

Ter Haar Romeny BM, van der Gon JJD, Gielen CCAM (1982) Changes in recruitment order of motor units in the human biceps muscle. Experimental Neurology 78:360–368.

Thomas CK, Ross BH, Calancie B (1987) Human motor-unit recruitment during isometric contractions and repeated dynamic movements. Journal of Neurophysiology 57:311–324.

Thomas CK, Ross BH, Stein RB (1986) Motor-unit recruitment in human first dorsal interosseous muscle for static contractions in three different directions. Journal of Neurophysiology 55:1017–1029.

Whitford M, Kukulka CG (2011) Task-related variations in the surface EMG of the human first dorsal interosseous muscle. Experimental Brain Research 215:101–113.

Zhou P, Suresh NL, Rymer WZ (2011) Surface electromyogram analysis of the direction of isometric torque generation by the first dorsal interosseous muscle. Journal of Neural Engineering 8:036028.

