## Supplementary figures and images for "Task-Dependent Motor Unit Recruitment and Rate Coding Reveal Redistribution of Neural Drive in the Human Hand"

### Supplementary Figure 1

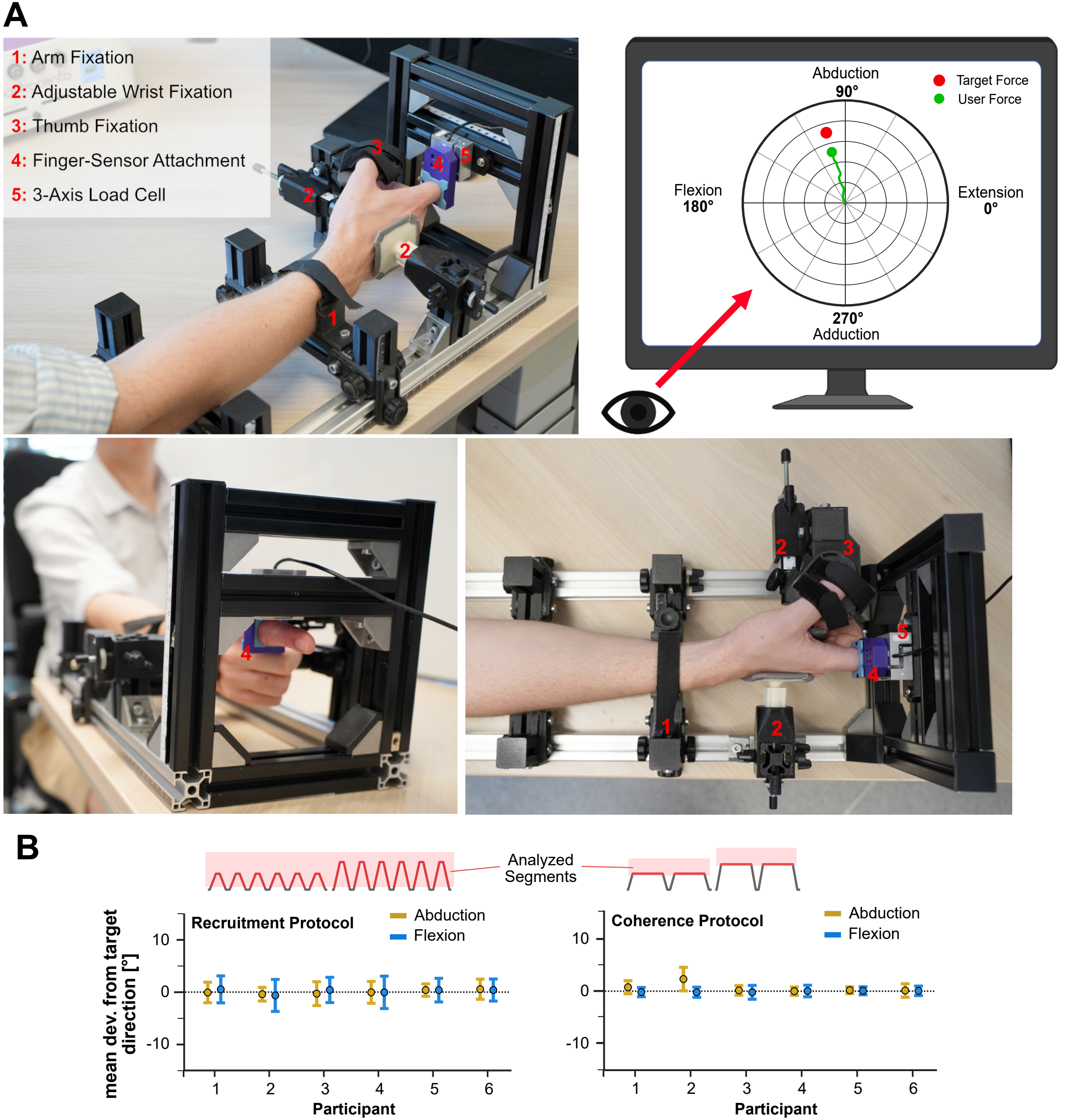

### Supplementary Figure 2

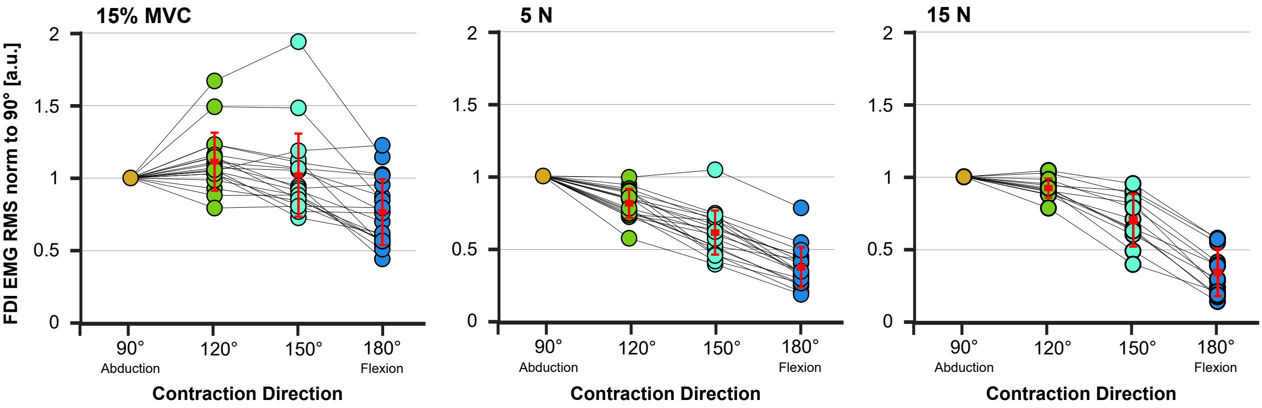

### Supplementary Figure 3

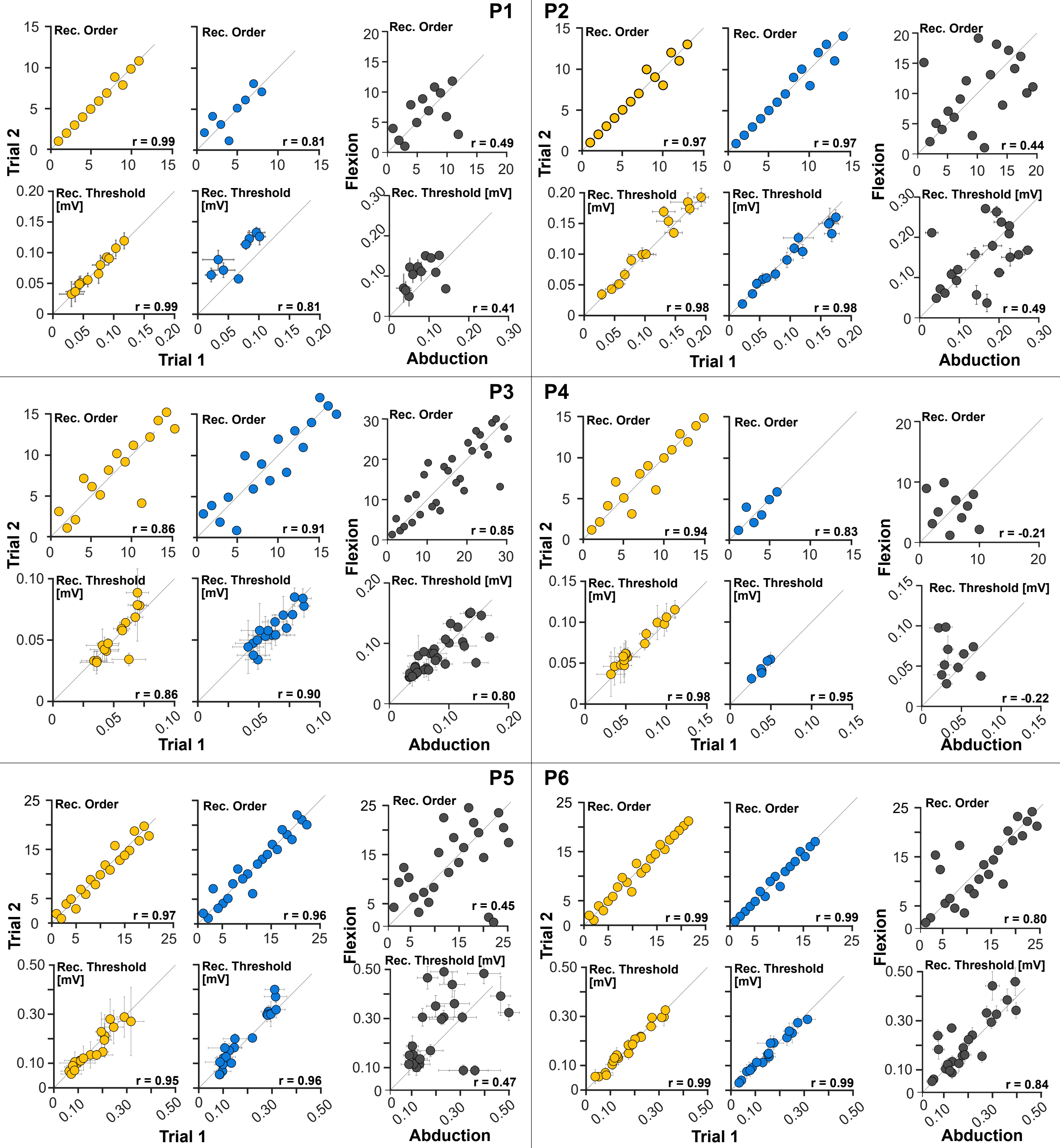

### Supplementary Figure 4

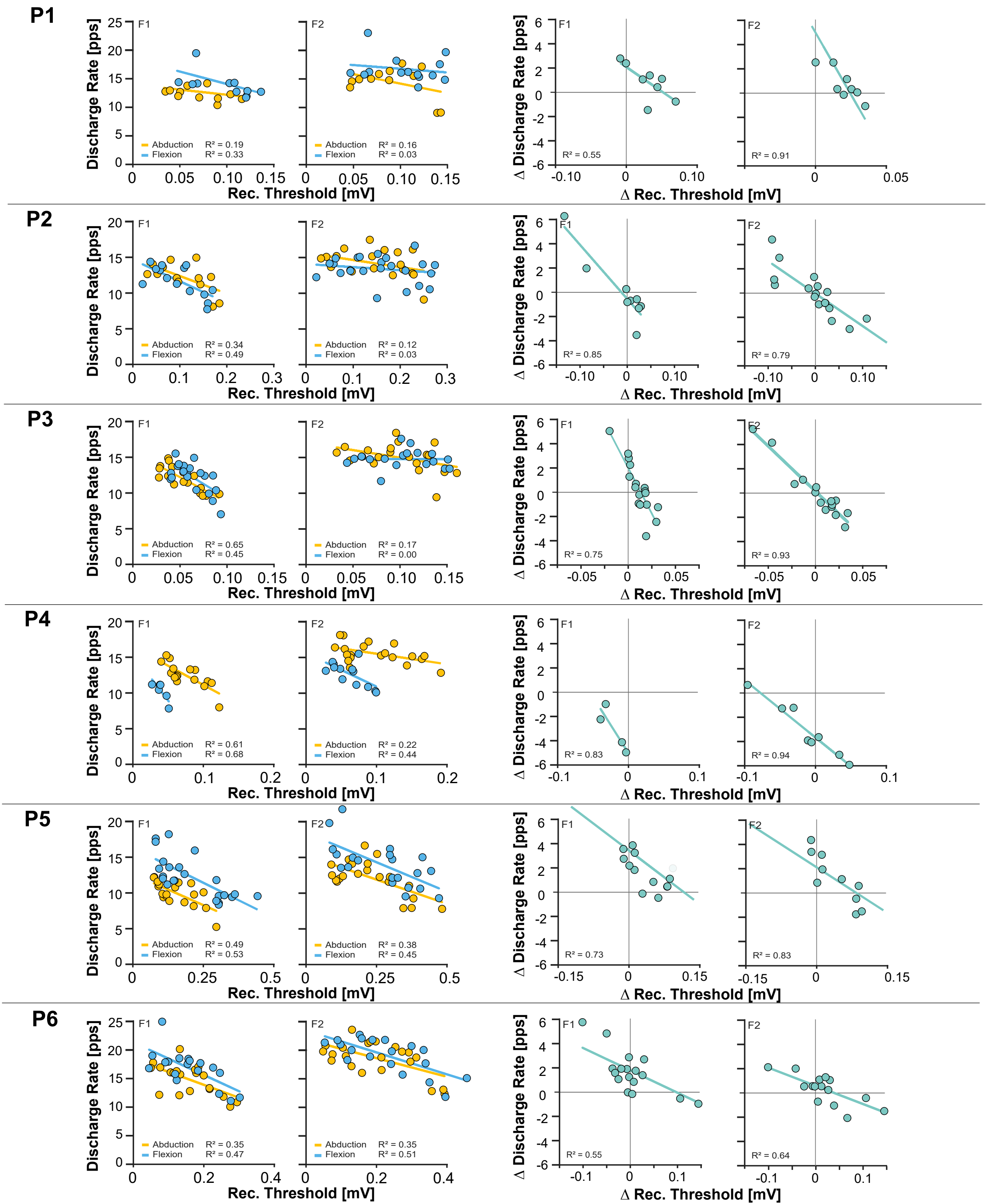
